# Application of Polymersomes Engineered to Target P32 Protein for Detection of Small Breast Tumors in Mice

**DOI:** 10.1101/187716

**Authors:** Lorena Simón-Gracia, Pablo Scodeller, Sergio Salazar Fuentes, Vanessa Gómez Vallejo, Xabier Ríos, Eneko San Sebastián, Valeria Sidorenko, Desirè Di Silvio, Meina Suck, Federica De Lorenzi, Larissa Yokota Rizzo, Saskia von Stillfried, Kalle Kilk, Twan Lammers, Sergio E Moya, Tambet Teesalu

## Abstract

Triple negative breast cancer (TNBC) is the deadliest form of breast cancer and its successful treatment critically depends on early diagnosis and therapy. The multi-compartment protein p32 is overexpressed and present at cell surfaces in a variety of tumors, including TNBC, specifically in the malignant cells and endothelial cells, and in macrophages localized in hypoxic areas of the tumor. Herein we used polyethylene glycol-polycaprolactone polymersomes that were affinity targeted with the p32-binding tumor penetrating peptide LinTT1 (AKRGARSTA) for imaging of TNBC lesions. A tyrosine residue was added to the peptide to allow for ^124^I labeling and PET imaging. In a TNBC model in mice, systemic LinTT1-targeted polymersomes accumulated in early tumor lesions more than twice as efficiently as untargeted polymersomes with up to 20% ID/cc at 24 h after administration. The PET-imaging was very sensitive, allowing detection of tumors as small as ∼20mm^3^. Confocal imaging of tumor tissue sections revealed a high degree of vascular exit and stromal penetration of LinTT1-targeted polymersomes and co-localization with tumor-associated macrophages. Our studies show that systemic LinTT1-targeted polymersomes can be potentially used for precision-guided tumor imaging and treatment of TNBC.

## Introduction

TNBC is defined by the lack of expression of estrogen receptor, progesterone receptor, and human epidermal growth factor 2. TNBC accounts for ∼15% of all breast cancer cases and is a heterogeneous group of tumors that can be classified in different subtypes based on gene expression profiles[1–3]. Some TNBC subtypes respond to the chemotherapy and have favorable prognosis, whereas other TNBC subtypes are aggressive with average cancer recurrence within 3 years after initial diagnosis and a life expectancy of ∼5 years[4]. These aggressive TNBC are locally invasive, highly metastatic, and must be detected and treated early to prevent dissemination.

Nanoformulations offer unique advantages for drug delivery. Nanoparticles can be designed to encapsulate hydrophobic molecules that would otherwise be insoluble, and payloads that have short circulation half-life and/or need to be protected from enzymes in the bloodstream, such as esterases or nucleases[5]. Cancer diagnosis and treatment can be combined into one modality by dual-use “theranostic” nanocarriers engineered to simultaneously deliver therapeutic and imaging cargoes[6][7]. Imaging payloads, such as fluorescent, MRI, and radio tags can be loaded in the nanosystems or coated on their surface. Nanosized polymeric vesicles (polymersomes) self-assembled from biocompatible copolymers are particularly appealing because of their versatility and unique properties. The high molecular weight of block copolymers results in the formation of highly entangled membranes displaying a high degree of resilience with elastomer-like mechanical properties. This confers the polymersomes a higher flexibility[8][9] and higher ability for tissue penetration than other vesicles self-assembled from low molecular weight entities, such as liposomes[10]. Polymersomes can be loaded with hydrophilic effector molecules, e.g. low molecular weight drugs[11][12], proteins[13], nucleic acids[14], and imaging agents[15][16], in their aqueous lumen and with hydrophobic cargoes within the polymer membrane[11][17].

The surface of nanoparticles can also be modified to improve their *in vivo* behavior such as circulation half-life, non-specific interactions and affinity for non-target sites, and to achieve selective accumulation in target tissue(s). Affinity ligands, such as peptides[18][19] and antibodies[20] can be coated on the nanoparticles for specific tissue and cell recognition. Tumor-penetrating peptides[21] can be used to concentrate cytotoxic molecules and drug-loaded nanoparticles in tumors and potentiate their antitumor activity[17][22]. The AKRGARSTA peptide, referred to as “LinTT1” (linear TT1), is a 9-amino acid tumor-penetrating peptide that binds to p32 protein. The primary receptor for LinTT1, p32, is a mitochondrial chaperone that is aberrantly expressed at the cell surface of malignant cells and activated tumor macrophages, which makes it a good target molecule for affinity-based cancer delivery[23][24]. A variety of solid tumors, such as glioblastoma and carcinomas of the gastrointestinal tract[24][25][18], overexpress cell surface p32 and several studies have found upregulated p32 expression in TNBC[24][26][27]. LinTT1 is processed by tumor-derived proteases, such as uPA, to C-terminally expose the C-end rule motif of the peptide (i.e. AKRGAR), which is capable of interacting with the cell- and tissue-penetration receptor NRP-1[23][28]. Recently, LinTT1-functionalization was found to significantly improve the therapeutic index of iron oxide nanoworms loaded with proapoptotic effector peptide in a model of peritoneal carcinomatosis[18] and also in a TNBC model in mice[27]. In that last study, the tumor accumulation of fluorescently labeled Lin-TT1 nanoparticles was evaluated by optical imaging of tissue sections. However, fluorescence imaging-based *in vivo* biodistribution studies remain challenging due to issues related to the low depth of light penetration, tissue autofluorescence, and the semi-quantitative nature of optical imaging[29].

PET and SPECT are clinically used for imaging of radioactive contrast agents with beta and gamma emission, respectively. As the signal only comes from the radiotracer and as the tissues do not possess inherent radioactivity, PET and SPECT are not subject to endogenous tissue background, as opposed to MRI, CT, and imaging using optical contrast agents [30]. Moreover, in PET and SPECT the signal is not affected by tissue depth (as during in vivo imaging using fluorescent tags) or respiratory motion (as in the case in MRI)[30].

Encouraged by the anticancer activity of LinTT1-targeted therapeutic nanoparticles on breast tumors in mice[27], we decided to evaluate polymersomes guided with the LinTT1 peptide as a potential theranostic nanocarrier to early detect TNBC lesions. We radiolabeled LinTT1-targeted PEG-PCL polymersomes and studied, for the first time, the homing to orthotopic small breast tumors and the biodistribution of the polymersomes using PET imaging in mice. Intravenously-administered p32-targeted ^124^I labeled polymersomes showed good tumor selectivity and, importantly, allowed detection of tumors smaller than 20mm^3^. Our results suggest potential applications of LinTT1 engineered polymersomes for early detection of TNBC.

## Results

### Preparation, functionalization and radiolabeling of polymersomes

Polymersomes were prepared by the film hydration method and functionalized with the Cys-Tyr-LinTT1 or Cys-FAM-LinTT1 peptide through a thioether bond between the maleimide group of the copolymer and the thiol group of the cysteine of the peptide. The number of polymersomes was determined by the ZetaView instrument and the functionalization with FAM-labeled peptide was quantified by fluorimetry. The peptide functionalization resulted in ∼3.7×10^4^ peptides/polymersome particle (density ∼0.7 peptides/nm^2^). For radiolabeling, the polymersomes were functionalized with the Cys-Tyr-Ahx-LinTT1 (Ahx = aminohexanoic acid) peptide or control Cys-Tyr dipeptide. The tyrosine residue was incorporated for radioiodination. The hydrodynamic diameter of LinTT1-Tyr-polymersomes (LinTT1-Tyr-PS), Tyr-polymersomes (Tyr-PS), polymersomes labeled with ATTO550 (LinTT1-ATTO550-PS and ATTO550-PS), and polymersomes labeled with FAM (LinTT1-FAM-PS and FAM-PS) measured by DLS, was ∼130nm for all the polymersome samples (Figure 1B). As shown in Fig. 1A, the size of polymersomes measured by TEM was smaller than 100nm. By TEM, we obtained the size of the dry polymersomes while DLS showed the hydrodynamic diameter of the particles in solution. In DLS the observed particle size was higher than in TEM due to DLS sample being in the solvated state with water molecules and ions associated with polymersomes[31]. Moreover, in the DLS, we used the intensity of the scattered light as a function of the particle size. As the intensity scales with the sixth power of the radius the larger particles have higher representation in the size distribution graphic. The Z-potential was slightly negative but very close to 0 for the different polymersome preparations (Figure 1B and S1). For the Z-potential measurements, we used a moderate ionic strength, 10mM. At high ionic strength, such as in phosphate saline buffer, the charges of the nanoparticles are screened by the ions in solution. At lower ionic strength, such as 10 mM NaCl, this effect is lower and the zeta potential is closer to the potential resulting from the actual charge of particle[32]. Moreover, by using a known ionic strength, the influence of unknown concentration of ions contained in water was avoided.

**Figure 1.**
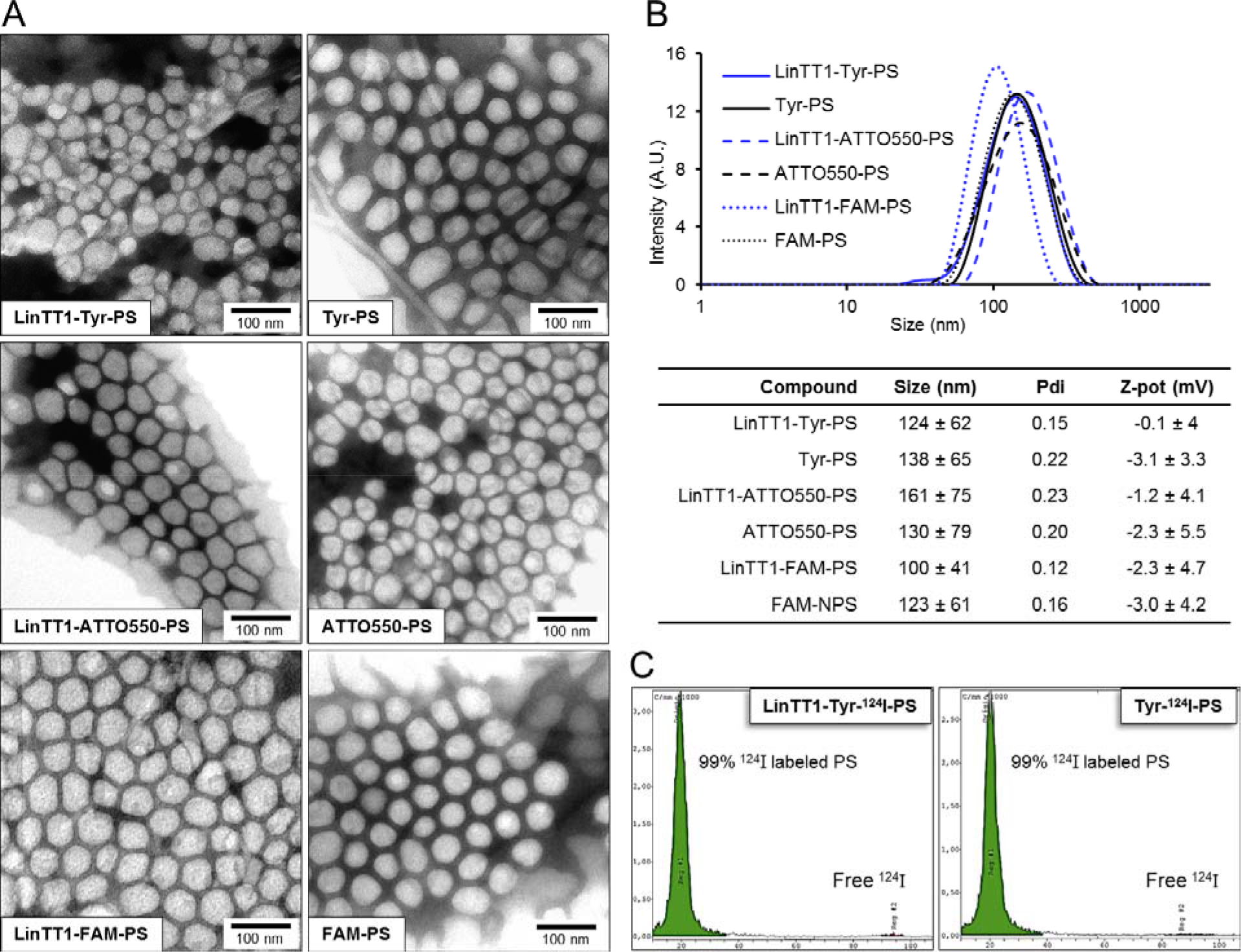
Characterization of the polymersomes. A) TEM images of the LinTT1-targeted and non-targeted radiolabeled polymersomes (LinTT1-Tyr-124I-PS and Tyr-124I-PS) and fluorescently labeled polymersomes (LinTT1-ATTO550-PS, ATTO550-PS, LinTT1-FAM-PS, and FAM-PS). B) Size distribution measured by DLS and summary of the physical properties of the polymersome preparations (3 independent measurements). C) Chromatograms obtained by TLC of radiolabeled polymersomes after the purification showing the percentage of ^124^I-labeled polymersomes and the peak of free ^124^I.

For PET imaging, the LinTT1-Tyr-PS and Tyr-PS were radiolabeled with ^124^I. Before purification, the efficiency of polymersome radiolabeling was determined by TLC (thin layer chromatography) (Figure S2). The yield of radiolabeling after purification, measured with activimeter, was 48±9% for LinTT1-Tyr-^124^I-PS and 43±2% for Tyr-^124^I-PS. The low radiolabeling of PEG-PCL polymersomes without peptide (14% or radiolabeling, Supplementary Figure 2) indicated that ^124^I present in LinTT1-Tyr-^124^I-PS and Tyr-^124^I-PS preparations was predominantly due to the covalent binding of ^124^I to the tyrosine residue of the peptides. TLC analysis after purification demonstrated that 99% of the ^124^I was bound to polymersomes (Fig. 1C).

### LinTT1-targeted polymersomes bind to recombinant p32 and to cultured breast tumor cells

To evaluate the effect of LinTT1 functionalization on the tropism of polymersomes *in vitro*, we first tested the binding of LinTT1-Tyr-^124^I-PS to human recombinant p32 protein, the primary receptor of LinTT1. P32-coated magnetic beads were incubated with the polymersomes, and polymersome binding was quantified by gamma counter. Compared to non-targeted polymersomes, LinTT1-Tyr-^124^I-PS showed ∼10-fold increased binding to the p32 beads (Figure 2A). This binding was specific, as the LinTT1-Tyr-^124^I-PS did not bind to NRP-1 (Figure 2A). The LinTT1 peptide does not bind to NRP-1 unless proteolytically processed by uPA[28]. These data show that the LinTT1 peptide attached to the polymersomes remains available for human p32 binding to modulate polymersome tropism.

**Figure 2.**
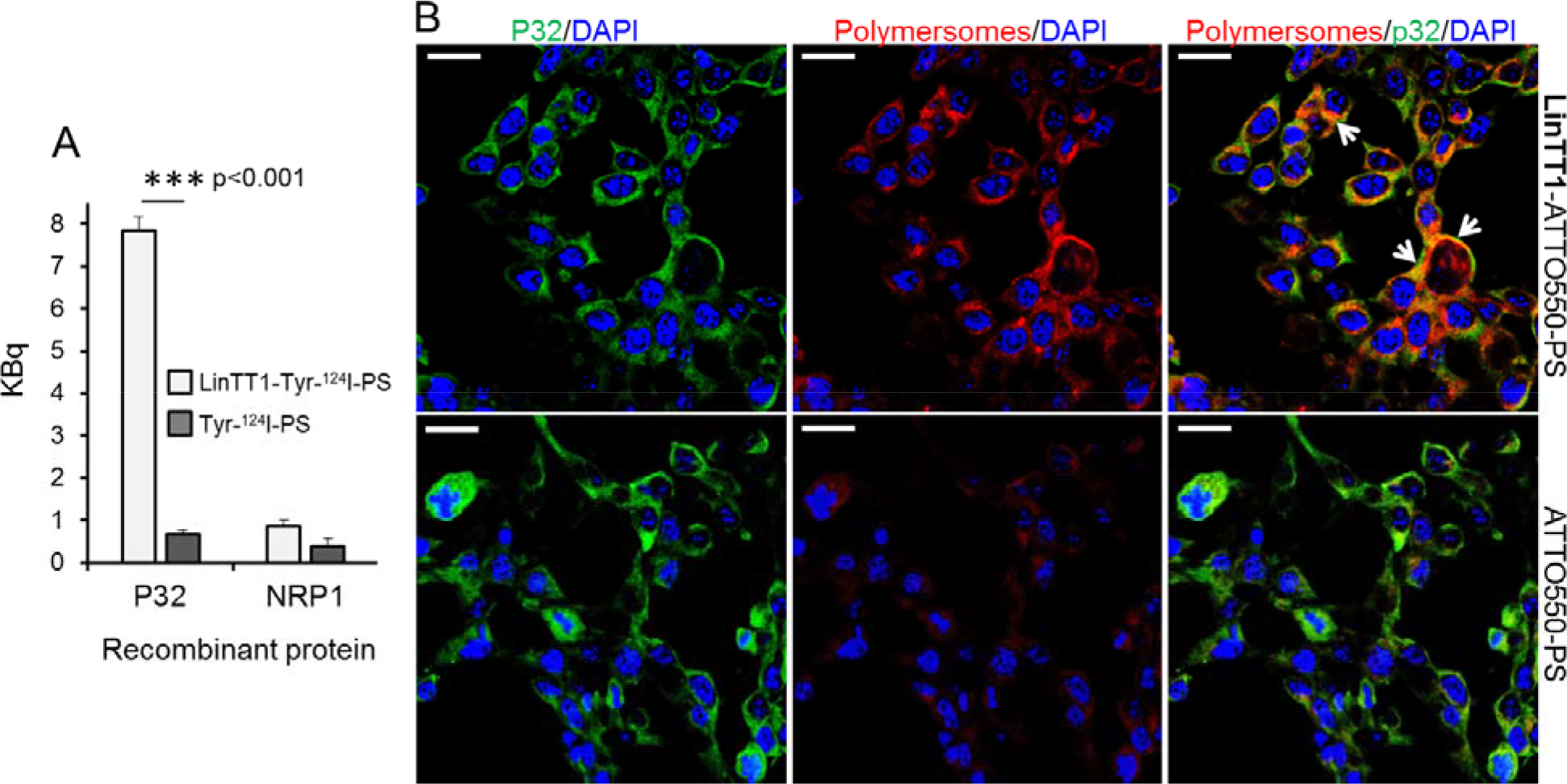
Binding of LinTT1-PS to recombinant p32 protein and to cultured 4T1 breast tumor cells. A) Binding of the LinTT1-Tyr-^124^I-PS and Tyr-^124^I-PS to p32 and NRP1-coated magnetic beads. The binding to the proteins after 1 h of polymersome incubation is expressed in KBq. N=3. Error bar=+SEM. B) Fluorescence confocal microscopy images of 4T1 cells incubated with LinTT1-ATTO550-PS or non-targeted ATTO550-PS for 1 h. The polymersomes were labeled with ATTO550 (red) and cells were immunostained for p32 protein (green). The nuclei were counterstained with DAPI (blue). Scale bar=20μm. White arrows point to the areas of colocalization of LinTT1-ATTO550-PS with p32.

Various human and mouse tumor cell lines express p32 on the cell surface[24]. We studied the presence of cell surface p32 in 4T1 and MCF-10CA1a TNBC cells by flow cytometry and confocal microscopy, and confirmed its surface expression on both cell lines (Figure S3). We tested the uptake of polymersomes and p32 co-localization in 4T1 cells - the same cell line used to establish the in vivo TNBC breast cancer model for systemic targeting studies. 4T1 cell line is of mouse origin and the tumors can be induced in immunocompetent Balb/c mice. The use of immunocompetent mice is an important aspect that is particularly relevant for the follow-up studies with LinTT1-guided combinations of companion diagnostic and therapeutic polymersomes. To study the uptake of polymersomes in 4T1 cells, we incubated the cultured cells for 1 h with LinTT1-targeted or control polymersomes labeled with ATTO550 (LinTT1-ATTO550-PS and ATTO550-PS) (Figure 2B). The LinTT1-functionalization increased polymersome uptake in 4T1 cells and the signal from LinTT1-ATTO550-PS partially colocalized with p32 (Figure 2B). We also studied the uptake of LinTT1-PS in MCF10CA1a human breast tumor cultured cells. Figure S4 showed a significantly higher uptake of LinTT1-PS compared with the non-targeted polymersomes. These experiments demonstrate that LinTT1 functionalization results in p32-enhanced uptake of polymersomes in cultured breast tumor cells.

### Systemic LinTT1 targeted radiolabeled polymersomes home to breast tumors

We used PET imaging to study *in vivo* biodistribution and tumor accumulation of systemic LinTT1-targeted polymersomes. Radiolabeled LinTT1-Tyr-^124^I-PS and Tyr-^124^I-PS were i.v. injected into mice bearing orthotopic 4T1 breast tumors and PET-CT scans were acquired at 10 min, 2, 6, 12, 24, and 48 h post-injection. To test whether detection of incipient breast tumors could be improved by targeting p32, the polymersomes were administered when breast tumor had reached ∼20mm^3^ (Figure 3).

**Figure 3.**
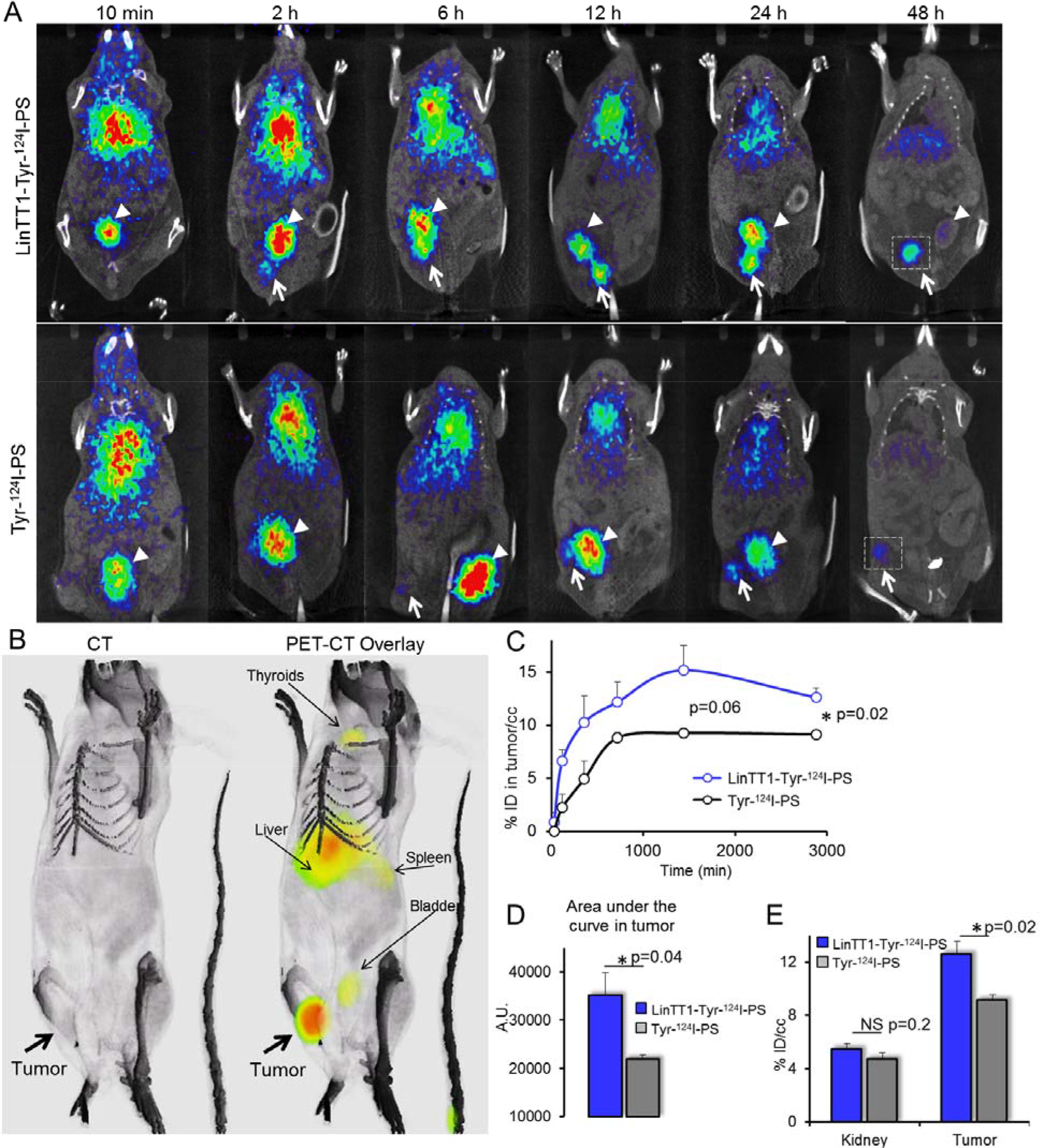
Radiolabeled LinTT1-PS home to 4T1 breast tumors. A) PET-CT imaging of 4T1 tumor mice injected with LinTT1-Tyr-^124^I-PS or non-targeted Tyr-^124^I-PS. White arrows point to the tumor. White arrowheads point to the bladder. The same mouse was used for the imaging at all the time points. The difference in tumor location is due to slightly different positioning of the mouse on the imaging support. B) 3D reconstruction of CT and PET-CT overlay images of mouse at 48 h after LinTT1-Tyr-^124^I-PS i.v. injection. C) Accumulation of radiolabeled polymersomes in the tumor. The % of ID/cc tumor was plotted against the time post-injection. The signal was quantified from the PET images. D) AUC of LinTT1-Tyr-^124^I-PS and non-targeted Tyr-^124^I-PS calculated from graph C. E) % ID/cc in kidney and tumor after 48 h of polymersome injection. The signal was quantified from the PET images. N=5 mice. Error bar=+SEM.

LinTT1 functionalization increased tumor homing of polymersomes at both early and late time points (Figure 3C), with the AUC (area under the curve) in tumor being ∼60% higher (Figure 3D). We saw tumor accumulation of LinTT1-Tyr-^124^I-PS already at 2 h post injection, whereas the tumor PET signal for non-targeted Tyr-^124^I-PS was only detectable at later time points

(Figure 3A). The highest tumor accumulation of LinTT1-Tyr-^124^I-PS was seen at 24 h after the injection, and it was 67% higher than for the untargeted polymersomes. At 48 h both targeted and untargeted polymersomes showed accumulation in the tumor (Figure 3A,B,E). At 48 h tumor accumulation of LinTT1-Tyr-^124^I-PS was lower than at 24 h, however, it was significantly higher than Tyr-^124^I-PS (12±0.9 and 9±0.4 ID/cc, respectively) (Figure 3C). In contrast, at 48 h, the signal in the kidney and thyroid gland in mice injected with targeted and untargeted polymersomes was not significantly different (Figure 3E, Figure S5).

The scans acquired at 6 h showed uptake of both targeted and non-targeted polymersomes in the liver (Figure 3A), in line with known role of the RES in the clearance of the circulating nanoparticles.

At 48 h after the injection, the tumors and organs were excised and ^124^I in tissue extracts was quantified with gamma counter. The highest percentage of ID/g of both targeted and nontargeted polymersomes after 48 h was observed in spleen and tumor (Figure 4A). Accumulation of both LinTT1 and untargeted polymersomes in spleen is consistent with the polymersome clearance by the RES. At 48 h, untargeted polymersomes showed accumulation in tumors (15±0.6% ID/g) and the functionalization with LinTT1 increased tumor accumulation of polymersomes by >70%, to 26±3% ID/g. Moreover, the percentage of ID/g of LinTT1-Tyr-^124^I-PS in tumor was 2.5 times higher than in liver (Figure 4A). Quantification of radioactivity revealed more than 2-fold higher accumulation of LinTT1-Tyr-^124^I-PS than Tyr-^124^I-PS in the sentinel lymph node of breast tumor mice (Figure 4A).

**Figure 4.**
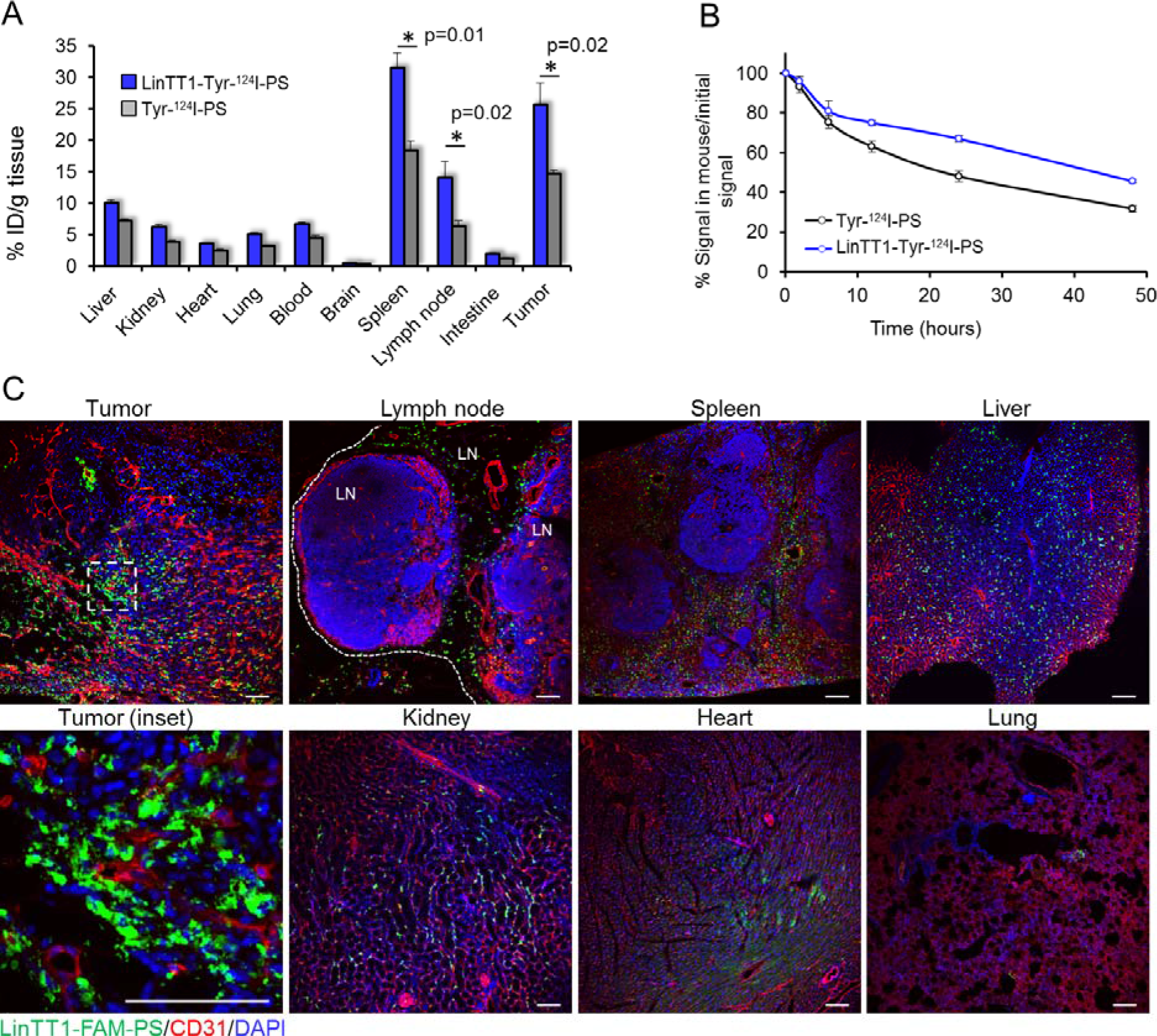
Biodistribution of radioactive and fluorescent polymersomes in 4T1 tumor mice. A) Biodistribution of i.v.-injected ^124^I labeled polymersomes in tumors and organs at 48 h after injection. Tumors, control organs, and blood were collected at 48 h post injection of radiolabeled polymersomes and the radioactivity was measured by gamma counter. N=6. Error bar=+SEM. B) Elimination rate of ^124^I quantified from the PET data. The radioactive signal of the whole mouse was determined at different time points. N=5 mice. Error bar=+SEM. C) Confocal fluorescence imaging of sections of 4T1 tumors and control organs from mice injected i.v. with LinTT1-FAM-polymersomes. Tissues were collected at 24 h post-injection of polymersomes into 4T1 bearing mice and sectioned and immunostained for FAM and CD31. Green: LinTT1-FAM-PS; red: CD31; blue: DAPI nuclear staining. LN= lymph node.

The elimination rate of ^124^I was studied by quantification of the PET imaging data. At 24 h, ∼50% of the injected Tyr-^124^I-PS and 67% of LinTT1-Tyr-^124^I-PS remained in the body. After 48 h, 32% of Tyr-^124^I-PS and 45% for LinTT1-Tyr-^124^I-PS remained in the body (Figure 4B). We have shown in a recent publication that an insignificant portion of the peptide is released from PEG-PCL polymersomes incubated with the serum of the 4T1 tumor bearing mice for 6 h[33]. We suggest that the high excretion at short time points observed is due to the renal clearance of the ^124^I released from the peptide-conjugated polymersomes. It is important to note that the signal in thyroid gland (Figure S5) - which accumulates free iodine - is similar for both targeted and untargeted polymersomes, suggesting similar leaching of iodine from polymersomes.

The effect of LinTT1 functionalization on the biodistribution and elimination rate of the polymersomes may be due to depletion of circulating LinTT1-polymersomes by the target sites: tumor tissue and macrophages. 4T1 tumor mice injected with both LinTT1 targeted and untargeted polymersomes showed similar ^124^I excretion rate at short time points. However, after 6 h, the ^124^I excretion rates for the targeted polymersomes became lower, likely due to preferential uptake of LinTT1-Tyr-^124^I-PS by p32+ tumor cells and activated macrophages.

### LinTT1-polymersomes target both the tumor cells and tumor macrophages

We next studied the tissue biodistribution of i.v. administered FAM-labeled polymersomes in 4T1 orthotopic tumor mice at the cellular level. The polymersomes were injected in 4T1 tumor mice, allowed to circulate for 24 h, and the sections of tumors and control organs were analyzed by confocal immunoanalysis.

We first studied the biodistribution of p32 immunoreactivity in tissues. In a previous report, p32 was found to be upregulated in MDA-MB-435 breast tumors compared to the control organs[34][24]. P32 immunostaining of sections of tumors and control organs from 4T1 mice demonstrated elevated expression of p32 in tumor tissue (Figure S6).

In agreement with the tissue extract-based radiography data, FAM-LinTT1-polymersomes accumulated in tumor and spleen (Fig. 3A). It was recently published that LinTT1 functionalization of nanoparticles enhances their penetration into tumor tissue[27][18]. Here we show that at 24 h, the LinTT1-FAM-PS in tumors did not colocalize with CD31-positive blood vessels, confirming that polymersomes had extravasated and penetrated into tumor stroma (Figure 4C, tumor inset). It has been shown that p32 is expressed by CD11b positive macrophages[34] and that LinTT1-conjugated nanoparticles colocalized with C68-positive macrophages in the breast[27], gastric, and colon tumors[18]. To study the macrophage uptake of LinTT1-PS, sections of tumors and organs were immunostained with antibodies against CD68, CD11b, and CD206 markers. CD68 and CD11b are pan-macrophage markers that label normal macrophages (including macrophages in spleen, lung, and in Kupffer cells in liver[33]), and TAMs[35]. CD206 is a marker of pro-tumor M2 macrophages[36], which promote tumor growth[37].

We found that LinTT1-FAM-PS colocalized with CD68 (□50% of colocalization), and showed partial colocalization with CD11b and CD206 (9% and 21% of colocalization, respectively) in tumors, confirming the targeting of tumor-associated macrophages (Figure 5A and 5B). CD68-positive macrophages, extensively found in sentinel lymph node and spleen, and in liver (to a lesser extent), also showed colocalization with LinTT1-FAM-PS (Figure S7A and S7B), which might be one of the reasons for the higher accumulation of LinTT1-polymersomes in spleen and lymph node (Fig. 4A) compared with untargeted polymersomes.

**Figure 5.**
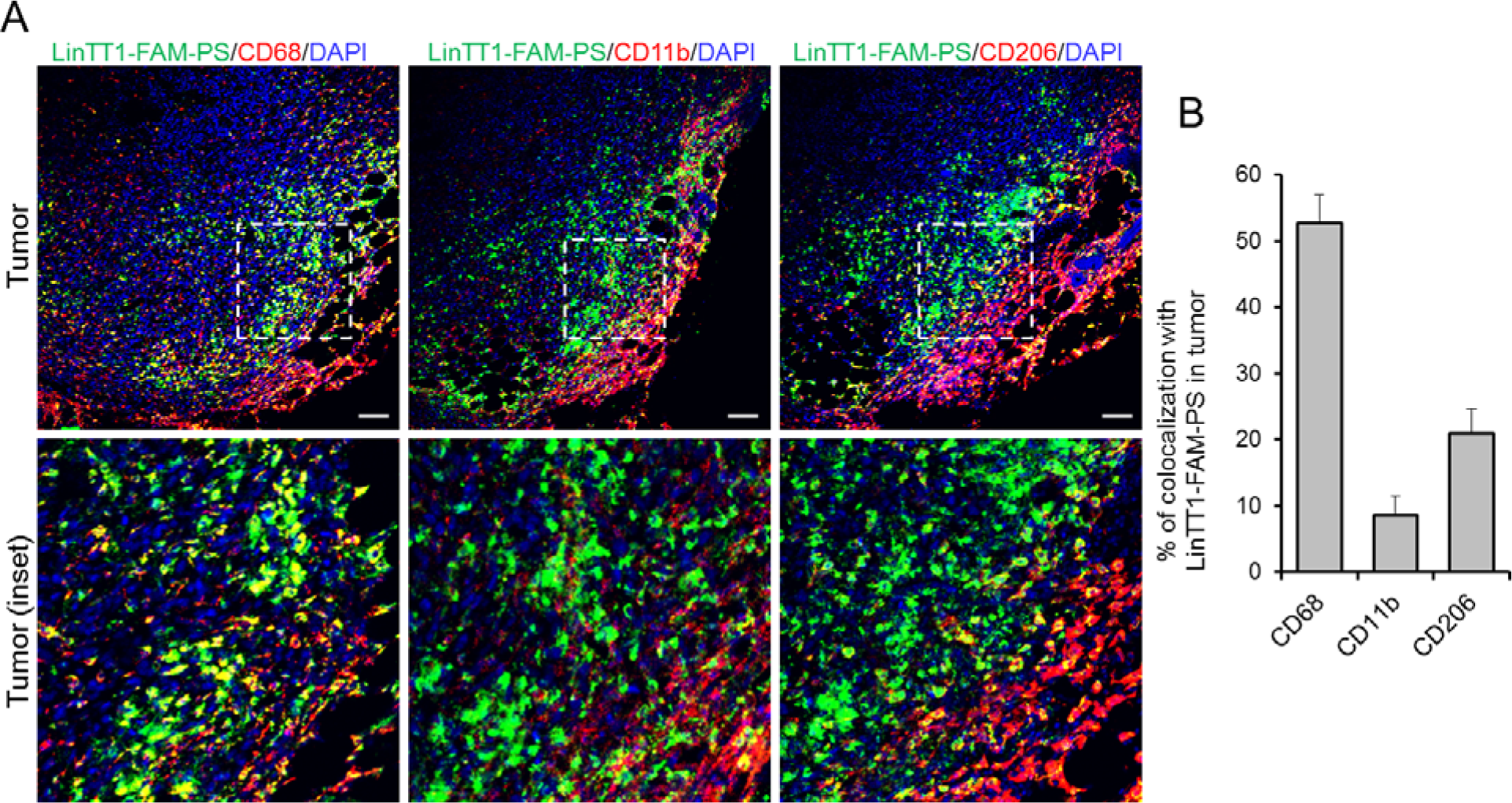
Colocalization of LinTT1-PS with macrophage markers in tumor tissue. 4T1 tumors were collected at 24 h after LinTT1-FAM-PS i.v. injection into 4T1 bearing mice, sectioned and immunostained. A) Confocal images of tumor sections immunostained for FAM, CD68, CD11b, and CD206, and counterstained with DAPI. Green: LinTT1-FAM-PS; red: CD68, CD11b, CD206; blue: DAPI counterstaining. Scale bar=100μm. B) Quantification of the colocalization of LinTT1-FAM-PS and macrophage markers in tumor using FLUOVIEW Viewer software. Error bar=+SEM.

### Human triple negative breast tumors and methastatic lymph nodes overexpress p32 and CD68+ macrophages

To further investigate the clinical relevance and translatability of LinTT1-targeted polymersomes, we investigated the distribution of p32 and CD68 immunoreactivity on surgical cases of triple negative breast primary tumors and metastasis, in comparison to healthy breast tissue. As exemplified in figure 6A and B, p32 diplays a uniform pattern of expression on healthy breast and healthy lymphoid tissue whereas strong membranous staining is found on malignant lesions. p32 is statistically significantly overexpressed in primary tumors, both metastatic and non-metastatic, and in particular on sentinel lymph nodes metastasis (Figure 6B,C).

**Figure 6:**
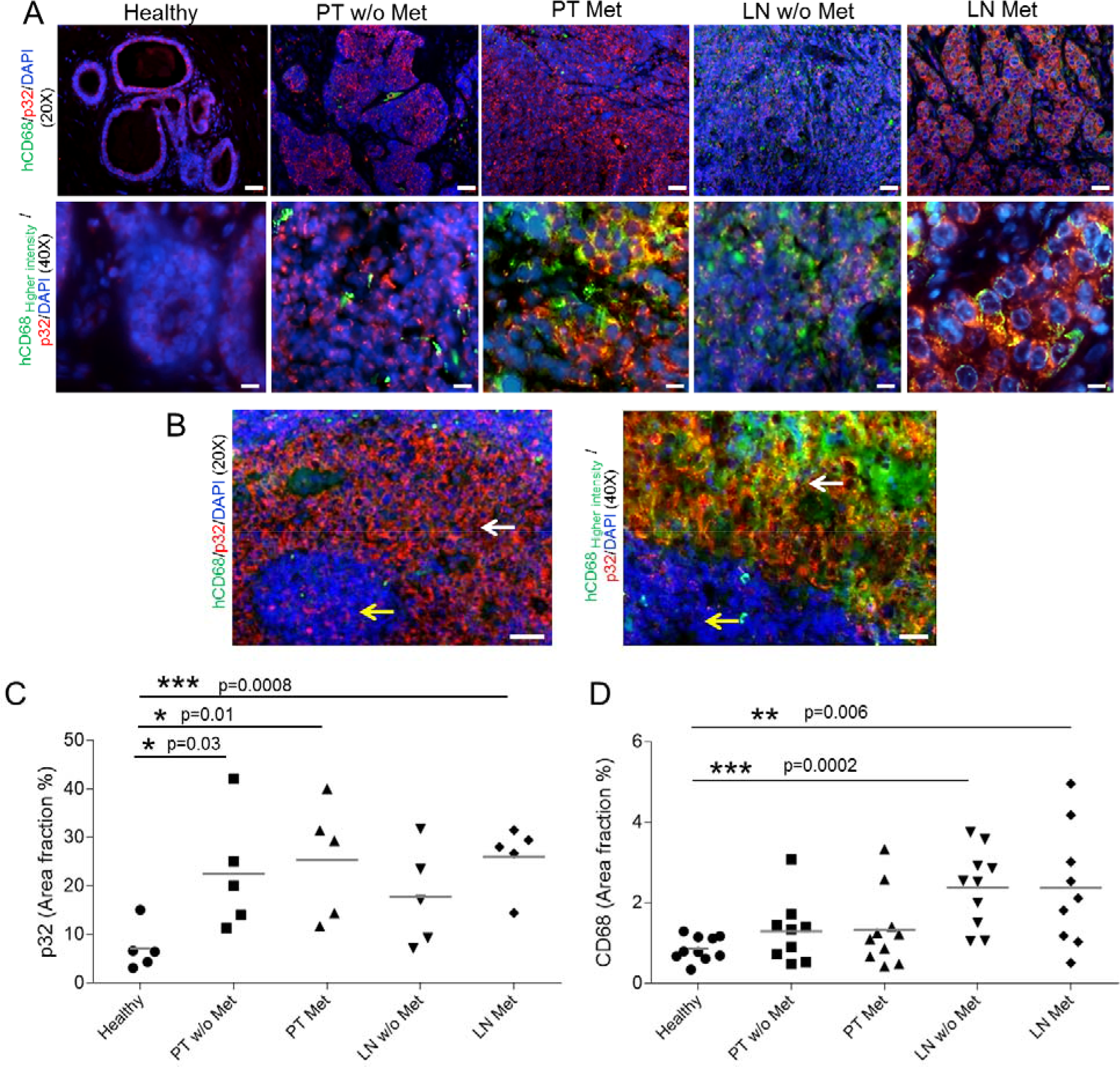
p32 and CD68 expression in human tissue samples. A) Immunofluorescence-based staining of human p32 and CD68 of FFPE human sections of healthy breast tissue (healthy), Primary Tumors with and without metastasis (PT Met and PT w/o Met, respectively) and their correspondent sentinel lymph nodes (LN Met and LN w/o Met, respectively), in N=5 patients/group. Representative pictures at 20x and 40x magnification. B) Transitioning areas in metastatic lymph node showing immunostaining in healthy tissue (yellow arrows) and malignant lesions (white arrows) in lymph node. The green intensity was increased to show the colocalization of p32 and CD68. C) Quantification of the p32 area fraction percentage in the different tissue sections. Three 20x representative images were acquired per patient and quantification was performed by the imaging-processing and analysis software AxioVision SE64 rel 4.8.3 (Zeiss Microscopy). N=5 patients/group. Scale bar for 20x=40 μm; scale bar for 40x=20 μm. D) Quantification of the CD68 in the tissue sections from immunohistochemistry analysis. Four 20x representative images were acquired per patient and quantification was performed by the Automated Training Segmentation algorithm from the InForm Software (Perkin Elmer). N=10 patients/group. Scale bar for 4x=400 μm; scale bar for 20x=100 μm.

Additionally, increased number of CD68-positive cells was found in breast tumors and in sentinel lymph nodes, both from patients with and without metastases, in comparison to healthy breast (Figure 6D and Figure S8). Partial colocalization of p32 and CD68 was also observed in malignant areas in metastatic lymph node (Fig. 6B), demonstrating the expression of p32 in activated macrophages.

## Discussion

In the current study, we evaluated the LinTT1-guided biocompatible PEG-PCL polymersomes as PET contrast agent for TNBC detection. Our findings indicate that LinTT1-polymersomes can be used for sensitive and specific detection of small triple negative breast tumors. This, along with recently published reports on LinTT1-mediated targeting of therapeutic nanocarriers[18][27], suggests potential theranostic applications for the LinTT1-targeted nanocarriers for early detection and treatment of TNBC.

Nanoparticles have been affinity targeted to tumors for PET imaging. For example, in a recent PET study, clinical application of RGD-targeted PET-active nanoparticles for melanoma imaging has been reported[38]. Systemic nanoparticles of ∼100nm are cleared by complement-mediated phagocytosis by Kupffer and endothelial cells of the liver as well as by phagocytic cells in the rest of the RES[39]. The polymersomes used in the current study are composed of biocompatible, biodegradable, and FDA approved PEG and PCL polymers[40]. In a dedicated toxicology study, the PEG-PCL-PEG micelles did not show signs of acute toxicity and there were no significant lesions found in histopathological study of major organs, including the liver and kidneys[41]. The maximum tolerated dose of these nanoparticles by i.v. administration was calculated to be 200-fold higher than the dose used in our study.

The current study documents high tumor accumulation of LinTT1-polymersomes (>20% ID/cc) that translates into an ability to detect very small malignant lesions, barely visible by CT. This sets our system apart from other molecular and nanoparticle PET contrast agents with reported tumor accumulations between 5-10% ID/g [42][43][44]. Remarkable tumor selectivity and tumor binding capacity observed for the LinTT1-polymersomes is likely to be due to a combination of the tumor homing properties of LinTT1 peptide with the favorable properties of the PEG-PCL polymersome nanoplatform. LinTT1 belongs to a family of tumor homing peptides that, unlike conventional vascular homing peptides, are not limited to vascular docking sites but have access to extended tumor extravascular space[45]. P32 protein, the receptor of LinTT1 peptide, is normally expressed in the mitochondria of the cells, but it is aberrantly displayed on the surface of tumor cells and on macrophage/myeloid cells, specially in hypoxic areas of tumors[23][24]. LinTT1 binds to the superficial p32 on the tumor cells and activated macrophages and is cleaved by uPA, an enzyme involved in tumor migration and progression[46]. LinTT1 then exposes the C-end motive (R/KXXR/K) on the C-terminal. C-end motif binds to NRP-1 protein, which is overexpressed in tumor cells and tumor vasculature. The binding to NRP-1 triggers an increase of the tumor tissue permeability and the peptide together with the cargo is internalized into the tumor[47]. Another potentially contributing aspect, not addressed in the current study, is the ability of LinTT1 to increase tumor penetration of co-administered compounds and nanocarriers. LinTT1-iron oxide nanoparticles were recently found to increase tumor penetration of co-administered 70kDa dextran[18]. Homing of LinTT1-nanocarriers may thus not be limited by the number of systemically accessible peptide receptors[48] and allow more nanocarriers to enter the tumor tissue for improved sensitivity of detection. Tumor accumulation of LinTT1-polymersomes may also be enhanced by physicochemical features of the PEG-PCL polymersomes used in the current study. On one hand, the flexibility of polymersomes[10] may contribute to tissue penetrative targeting with tumor-penetrating peptides. In addition, polymersomes are known to possess an intrinsic tumor tropism. The passive targeting of polymersomes presents an additional advantage for the tumor detection and treatment. We have recently demonstrated that pH-sensitive polymersomes efficiently delivered payloads to the tumor tissue in the absence of active targeting[11]; this accumulation was further boosted by targeting with iRGD peptide[17]. Likewise, the systemic radiolabeled non-targeted polymersomes in the current study showed high accumulation in 4T1 breast tumors; this accumulation was potentiated by functionalization of polymersomes with the LinTT1 peptide by about 70%. The effect of LinTT1 targeting is more prominent at earlier time points (at 2 h post-injection the LinTT1-PS, but not non-targeted PS, are already visible in the tumor, Fig. 3), a useful feature that can reduce the imaging acquisition time for the tumor detection.

LinTT1 tumor homing is likely to be due to a combination of targeting of tumor cells and tumor-associated macrophages (TAMs). The targeting of both these cell populations increases the accumulation of LinTT1-PS in tumor to facilitate tumor detection. In the 4T1 breast tumor mice, the highest ID/g of LinTT1-polymersomes was seen in the tumor, spleen, and lymph nodes. All these tissues contain abundant macrophages, a cell population known to upregulate, upon activation, the expression of cell surface p32. TAMs are an important therapeutic target, which play major roles in progression of solid tumors and can act as slow-release reservoir of drugs encapsulated in polymeric particles[49]. LinTT1-polymersomes may be capable of targeting tumor cells and TAMs in TNBC patients, as the peptide is not species specific, and since TAMs are abundant in clinical lymph node and breast tumor samples. We show here that primary breast tumors and sentinel lymph nodes from clinical samples from patients overexpress p32 protein and that the number of CD68+ macrophages is increased compared with healthy tissues. This finding supports potential translatability of the system into clinical applications for the detection and treatment of TNBC. Clinical breast tumors are heterogeneous and the cell surface p32 expression and the sensitivity to p32-targeting-based treatment is likely to differ between the patients. PET imaging with LinTT1-polymersomes can be potentially used as a companion diagnostic test for selection of patient cohort most likely to respond to p32-targeted therapies.

Lymphangiogenesis in tumor-draining lymph nodes occurs before the onset of metastasis and is associated with distant metastasis. A study using Lyp-1 (a tumor lymphatics-specific peptide and also p32-binder[24]) to image tumor-induced lymphangiogenesis[50], suggested that the pre-malignant niche is positive for p32. The accumulation of LinTT1-polymersomes in sentinel lymph nodes containing 4T1 tumor cells migrating out from the primary tumor and activated macrophages, suggests potential applications for LinTT1-polymersomes for more precise detection of early metastatic dissemination of breast cancer than is possible with currently approved compounds, such as Lymphoseek[51].

The potential applications of our system extend beyond breast cancer detection and therapeutic targeting. Systemically accessible p32 is overexpressed across solid tumors, including, gastric, colon, and ovarian caricinoma[18], glioma (Säälik et al. unpublished), and in atherosclerosis lesions[52]. Systematic evaluation of the relevance of the linTT1-polymersomes for detection and/or therapy of these conditions will be a subject of follow-up studies.

## Materials and Methods

### Materials

Polyethylene glycol-polycaprolacone (PEG_5000_-PCL_10000_, PEG-PCL) and Maleimide-PEG_5000_-PCL_10000_ (Mal-PEG-PCL) copolymers were purchased from Advanced Polymer Materials Inc. (Montreal, Canada). Cys-Tyr-Ahx-LinTT1 (Ahx = aminohexanoic acid) and Cys-Tyr peptides were purchased from KareBay Biochem, Inc. (USA), and Cys-fluorescein (FAM)-TT1 and Cys-FAM peptides were purchased from TAG Copenhagen (Denmark). Sodium iodine-124 was purchased from Perkin Elmer (Amsterdam). Thin liquid chromatography sheets were purchased from Agilent Technologies (USA). The ATTO550-amine dye was purchased from Atto-Tech GmbH (Germany). 4T1 cells were purchased from (ATCC, CRL-2539) were cultured in RPMI medium 1640 + GlutaMAX with 25mM HEPES (Gibco, Life Technologies, USA) containing 100IU/mL of penicillin and streptomycin, and 10% of FBS (GE Healthcare, UK). MCF10CA1a cells were obtained from Erkki Rouslahti (Cancer Research Center, Sanford Burnham Prebys Medical Discovery Institute). The cells were maintained in Dulbecco’s Modified Eagle Medium (DMEM, Lonza, Switzerland) with 4.5mg/mL of glucose, 100IU/mL of penicillin and streptomycin, and 10% of FBS.

### Synthesis and characterization of peptide-PEG-PCL vesicles

PEG-PCL (8mg, 0.53μmol) and Mal-PEG-PCL (2mg, 0.13μmol) copolymers were dissolved in 1mL of acetone previously purged with nitrogen. The solvent was evaporated and the polymer film was hydrated with 1mL of PBS 10mM pH 7.4, previously purged with nitrogen. The suspension was sonicated for 5 min and the peptide (Cys-Tyr-LinTT1, Cys-Tyr, Cys-FAM-LinTT1peptide, or Cys-FAM, 0.4mg, 2eq), dissolved in 0.2mL PBS previously purged was added to the suspension. The suspension was sonicated for 30 min, mixed at room temperature for 2 h and kept overnight at 4°C. The vesicles were purified using centrifugal filters of 100kDa MWCO (Amicon Ultra, Merck Millipore. Ltd. Ireland) and the final suspension was concentrated to 100mg of copolymer/mL.

For the polymersomes labeled with ATTO550, the dye was first conjugated to the polymer. Mal-PEG-PCL (10mg, 0.65μmol) was dissolved in 0.3mL of DMF previously purged with nitrogen and ATTO550-NH_2_ (0.77mg, 2eq) dissolved in 0.1mL of previously purged DMF was added to the solution. Triethylene amine (1μL) was added to the solution as a catalyzer. The solution was reacted at room temperature overnight, dialyzed against water using dialysis membrane of 3.5KDa MWCO (Sigma-Aldrich), and freeze-dried. ATTO550-PEG-PCL (1mg, 0.06μmol), PEG-PCL (7mg, 0.47μmol) and Mal-PEG-PCL (2mg, 0.13μmol) were dissolved in 0.5mL of acetone. The solvent was then evaporated to form the polymer film. The polymersomes were assembled and the Cys-LinTT1 peptide conjugation was performed as described above.

DLS and Z-potential measurements (Zetasizer Nano ZS, Malvern Instruments, USA) were used to assess the average size, polydispersity, and surface charge of polymersome preparations. The size was measured at a concentration of 1mg polymer/mL in PBS (10mM of phosphate and 137mM of NaCl). The z-potential was measured at 0.2mg of polymer/mL in NaCl 10mM). TEM was used to assess the size and morphology of assembled vesicles. Briefly, polymersomes were deposited from a water solution onto copper grids at 1mg/mL, stained with 0.75% phosphotungstic acid (pH 7), air-dried, and imaged by TEM (Tecnai 10, Philips, Netherlands). The number of polymersomes in the suspension was measured using the ZetaWiew instrument (Particle Metrix GmbH, Germany).

### Iodination of PEG-PCL vesicles

Two milligrams of Iodogen (Sigma-Aldrich, Spain) were dissolved in 10mL of CH_2_Cl_2_ and 20μL of this solution was transferred to a tube and the solvent was evaporated. LinTT1-Tyr-polymersomes or Tyr-polymersomes (1mg) were mixed with Na^124^I (18.5MBq) and 10µL of buffer phosphate 0.5M in a tube containing Iodogen. After 30 min 250μL of phosphate buffer, 1M NaCl, pH 7.4 was added to the reaction and the solution was transferred to a tube containing 50μL of Na_2_S_2_O_3_ 0.1M. The radiolabeling yield was measured by TLC using glass microfiber chromatography paper impregnated with silica gel (Agilent Technologies, USA) and ethanol:water 85:15 as mobile phase. The radioactivity of the peaks was measured with a TLC reader (γ-MiniGITA, Raytest, Germany). The polymersomes were purified using centrifugal filters of 100kDa MWCO (Amicon Ultra, Merck Millipore. Ltd. Ireland) and resuspended in 0.1mL of PBS. The removal of the free ^124^I was confirmed by radio-TCL and the final radioactivity was measured with a dose calibrator (Capintec CRC-25R, USA).

### *In vitro* binding of polymersomes to recombinant p32 protein

Recombinant hexahistidine–tagged p32 was bacterially expressed and purified as previously described[23]. For protein binding assays, Ni-NTA magnetic agarose beads (Qiagen, Germany) in binding buffer (50mM Tris pH 7.4, 150mM NaCl, 5mM imidazole) were coated with p32 protein at 15μg of protein/10μL beads. Radiolabeled polymersomes were incubated with the p32-coated beads in binding buffer containing 1% BSA at room temperature for 1 h. The magnetic beads were washed with binding buffer and resuspended in a final volume of 1mL of binding buffer. The radioactivity of each sample was quantified by automatic gamma counter (2470 Wizard 2, Perkin Elmer).

### Assessment of *in vitro* cell surface p32 expression by flow cytometry

4T1 and MCF10CA1a cells were detached from the culture plate with enzyme-free cell dissociation buffer (Gibco, Life Technologies, UK). The cells (10^5^ cells) were incubated with 10μg/mL of the in-house generated rabbit polyclonal p32 antibody in blocking buffer containing 1% of BSA, 1% FBS, and 1% of goat serum in PBS at room temperature for 1 h. The cells were washed, and incubated with Alexa 647-conjugated goat anti-rat IgG (1/1000, Invitrogen, Thermo Fisher Scientific, MA, USA) in blocking buffer at room temperature for 30 min. After washes, the cell surface p32 expression was analyzed by flow cytometry (Accuri, BD Biosciences, CA, USA).

### Assessment of *in vitro* cell surface p32 expression by immunostaining

4T1 and MCF10CA1a cells (10^4^) were seeded on glass coverslip in a 24 well-plate and incubated overnight at 37°C. The cells were blocked in PBS containing 0.05% Tween-20, 5% FBS, 5% BSA, and 5% goat serum (GE Healthcare, UK) for 1 h. The cells were immunostained with 10μg/mL of the in-house generated rabbit polyclonal p32 antibody in buffer containing 1% of BSA, 1% FBS, and 1% of goat serum in PBS at room temperature for 1 h. The cells were washed, and incubated with Alexa 647-conjugated goat anti-rat IgG (1/2000, Invitrogen, Thermo Fisher Scientific, MA, USA) in blocking buffer at room temperature for 30 min. Cells were counterstained with 1μg/mL of DAPI, transferred to glass slides, and examined by fluorescence confocal microscopy using the Zeiss LSM710 instrument.

### Uptake of polymersomes in cultured cells

4T1 cells (5×10^5^) were seeded on glass coverslips in a 24-well plate and the next day incubated with ATTO550-labeled polymersomes (0.5mg polymer/mL) at 37 °C for 1 h. Cells were washed with PBS, fixed with 4% paraformaldehyde, permeabilized with 0.5% saponin, and blocked for 1 h with 1% BSA, 1% goat serum, 0.3M of glycine, and 0.05% of Tween-20 in PBS. Cells were then stained for p32 protein using anti-p32 rabbit polyclonal antibody (1/500, Millipore, Germany) and Alexa Fluor 488 goat anti-rabbit IgG (1/1000, Abcam, UK) as a secondary antibody, and counterstained with 1μg/mL of DAPI. Cells were examined under the confocal laser scanning microscope (LSM 510 Meta, Zeiss, Germany) equipped with a 63x oil objective lens (1.4 NA). Images were acquired sequentially to avoid cross-talk using excitation wavelengths 405, 488, and 561nm. Transmission images were collected and overlaid by Zeiss Zen software.

### *In vivo* PET-CT imaging

For tumor induction, 1 million 4T1 cells in 50μL of PBS were orthotopically implanted in the mammary gland of Balb/c mice (Charles River Laboratories, Spain). After 3 days, when the tumor volume had reached ∼18 mm^3^, the mice were injected in tail vein with 3.7-7.4MBq of radiolabeled polymersomes (1mg polymer, 100μL, N=5 mice) and subjected to PET-CT scans. During the scan acquisitions the mice were kept anesthetized with 1.5–2% isofluorane blended with O_2_. The PET-CT scans were performed at 10 min, 2, 6, 12, 24, and 48 h using the Argus PET-CT scanner (Sedecal, Molecular Imaging, Spain). First, PET scans were acquired using the following acquisition protocol: whole body emission scan, 2 beds, 10 min of total acquisition time for the scans at 10 min, 2, 6, 12, and 24 h post-injection and 20 min for the 48 h time point. The acquisition method was static, using 400-700KeV energetic window, FBP reconstruction algorithm, with correction for scatter coincidences. For the CT scans the used current was 140μA, 40kV of voltage, rotation of 360 degrees, 4 shoots, 1 bed, acquisition time of 6 min, and a reconstruction binning of 2. After 48 h of radiolabeled polymersome injection the mice were sacrificed and the tumor, blood, and organs were excised and further used for biodistribution studies. The PET-CT images were processed with the Medical Image Data Examiner AMIDE software. The CT and PET images were overlaid, the tumor volume was manually extracted from the CT scans and the same ROI was applied in the PET images, averaged, and expressed as percentage of injected dose per cubic centimeter of tissue (ID/cc). For the rendered 3D PET-CT images, PMOD image analysis software (PMOD Technologies Ltd, Zürich, Switzerland) was used. A 3D Gauss Filter of 1.5×1.5×1.5mm was applied to the PET image in order to increase the signal to noise ratio for 3D visualization.

### Biodistribution studies

For biodistribution studies, the tumor, blood, and organs were weighed and the radioactivity was measured using the automatic gamma counter. A standard curve was generated using ^124^I to determine the relationship between cpm and Bq. The biodistribution was expressed as percentage of injected dose per gram of tissue (ID/g). To determine the elimination rate of polymersomes, the radioactive signal in the whole mouse body was measured from the PET images at 10 min, 2, 6, 12, 24, and 48 h post-injection and normalized by the signal at 10 min post-injection. The elimination rate was expressed as signal in mouse x 100 divided by the signal at time zero.

### Tissue immunofluorescence and confocal microscopy

Balb/c mice were orthotopically injected with 1 million of 4T1 cells in the mammary gland and after 3 days FAM-LinTT1-PS (1mg of polymer, 100μL) was intravenously injected. After 24 h, the animals were sacrificed and the tumor and organs were excised, fixed in 4% of paraformaldehyde, cryoprotected with 15% and 30% sucrose, frozen down with liquid nitrogen, and cryosectioned at 10μm. Tissue sections were permeabilized using PBS 10mM containing 0.2% Triton-X for 10 min, and blocked in PBS 10mM containing 0.05% Tween-20, 5% FBS, 5% BSA, and 5% goat serum (GE Healthcare, UK) for 1 h. The sections were immunostained at dilution 1/100 with anti-fluorescein rabbit IgG fraction (Thermo Fisher Scientific, MA, USA), rat anti-mouse CD31, biotin rat anti-mouse CD11b, (BD Biosciences, CA, USA), rat anti-mouse CD68, rat anti-mouse CD206 (Bio-Rad, USA), and anti-p32 rabbit polyclonal antibody (Millipore, Germany) as primary antibodies. As secondary antibodies, Alexa 488-conjugated goat anti-rabbit IgG and Alexa 647-conjugated goat anti-rat IgG (1/500, Invitrogen, Thermo Fisher Scientific, MA, USA) were used. The sections were counterstained with DAPI and examined by fluorescence confocal microscopy using Olympus FV1200MPE instrument. The images were processed and analyzed using the FV10-ASW 4.2 Viewer image software (Olympus, Germany) and the Image J software.

### Immunofluorescence staining and quantification of p32 and CD68 in human tissue of breast tumor, lymph node, and healthy tissue

Surgical samples of TNBC with lymph node metastasis (PT (Primary Tumor) Met; n=5), TNBC without lymph node metastasis (PT w/o Met; n=5) as well as their corresponding metastasized sentinel lymph nodes (LN Met; n=5) and non-metastasized lymph nodes (LN w/o Met; n=5) and healthy breast tissue (healthy; n=5), were fixed for 12-24 hours in 4% neutral buffered formalin. Samples were dehydrated, embedded in paraffin, sectioned at 4μm, and mounted on coated microscope slides (Dako, Denmark). After deparaffinization and rehydration of the sections, antigen retrieval was performed in citrate buffer at pH 6 followed by endogenous peroxidase blocking for 10 min. The samples were then incubated for 1 h with blocking solution containing 5% FBS, 5% BSA, 5% donkey serum, and 5% goat serum in PBS, followed by incubation with rabbit anti-p32 antibody (1:200, made in house) and mouse anti-human CD68 antibody (1:200, Dako) at room temperature for 2 h. Next, slides were incubated at room temperature for 1 h with the secondary antibodies Cy3-conjugated donkey anti-rabbit IgG (1:500, Dianova) and Alexa 488-conjugated goat anti-mouse IgG (1:50, Dianova). The sections were counterstained with DAPI and examined by fluorescence microscopy (Zeiss^®^ AxioImager M2 microscope). The quantification was performed by the imaging-processing and analysis software AxioVision SE64 rel 4.8.3. All stainings were performed on archived FFPE human samples and approved by the local ethics committee (EK 039/17).

### Immunohistochemistry staining and quantification of CD68 in human tissue of breast tumor, lymph node, and healthy tissue

Surgical samples of TNBC with lymph node metastasis (PT Met; n=10), TNBC without lymph node metastasis (PT w/o Met; n=10) as well as their corresponding metastasized sentinel lymph nodes (LN Met; n=10) and non-metastasized lymph nodes (LN w/o Met; n=10) and healthy breast tissue (healthy; n =10), were fixed for 12-24 hours in 4% neutral buffered formalin. In two cases with neoadjuvant chemotherapy and in two cases where only very small residual tumor was found in the surgical specimen, specimens from corresponding diagnostic biopsies were selected for immunohistochemistry. Samples were dehydrated, embedded in paraffin, sectioned at 4 μm, and mounted on coated microscope slides (Dako, Denmark). After deparaffinization and rehydration of the sections, antigen retrieval was performed in citrate buffer at pH 6 in a pre-treatment module (PT-Link, Dako, Denmark). Using an autostainer (Thermo Fisher Scientific) slides were incubated with endogenous peroxidase blocking solution for 5 min, followed by mouse anti-human CD68 antibody (1/100, Dako) for 30 min. Next, slides were incubated with secondary antibodies goat anti-mouse IgG conjugated to a peroxidase-labeled polymer chain (Dako, Denmark) for 20 min. The antigen-antibody-polymer complex was visualized with DAB + Chromogen (Dako, Denmark) for 10 min. The counterstaining was performed with Hematoxylin (Dako, Denmark) for 5 min. Finally, slides were covered with coverslipping film (Sakura, 6132 Prisma®) and examinated with Vectra^®^ 3 automated quantitative pathology imaging system (Perkin Elmer). Quantification was performed by the Automated Training Segmentation algorithm from the InForm Software (Perkin Elmer). All stainings were performed on archived FFPE human samples and approved by the local ethics committee (EK 039/17).

### Statistical analysis

All the statistical analysis was performed with the Statistica 8 software, using the one-way ANOVA, Fisher LSD test.

## Abbreviations

TNBC: triple negative breast cancer
PEG-PCL: polyethylene glycol-polycaprolactone
ID: injected dose
cc: cubic centimeter
MRI: magnetic resonance imaging
uPA: urokinase type plasminogen activator
NRP-1: neuropilin-1
PET: positron emission tomography
SPECT: single photon emission computed tomography
CT: computed tomography
FAM: fluorescein
DLS: dynamic light scattering
TLC: thin layer chromatography
i.v.: intravenously
AUC: area under the curve
RES: reticuloendothelial system
TAMs: tumor-associated macrophages
FFPE: formalin-fixed paraffin-embedded
FBS: fetal bovine serum
PBS: phosphate buffer saline
DMF: dimethyl formamide
TEM: transmission electron microscopy
BSA: bovine serum albumin.

## Author Contributions

### Lorena Simón-Gracia

Study conception, experiment design, polymersome synthesis, radiolabeling, PET-CT scan acquisition, flow cytometry, cells and tissue immunostaining, microscope pictures acquisition, data analysis, manuscript writing

### Pablo Scodeller

Study conception, experiment design, PET-CT scan acquisition, PET images analysis and quantification, cell immunostaining, microscope pictures acquisition, breast tumor induction, manuscript writing and revision

### Sergio Salazar Fuentes

Performance of animal experiments: breast tumor induction, animal manipulation during the PET-CT scans, tumor and organs excision

### Vanessa Gómez Vallejo

supervision of the radiolabeling

### Xabier Ríos

practical support for the radiolabeling

### Eneko San Sebastián

practical support for the PET-CT scans, PET-CT images analysis

### Valeria Sidorenko

polymersome uptake in cultured cells

### Desiré Di Silvio

practical support for the confocal microscope pictures acquisition

### Meina Suck, Federica De Lorenzi, Larissa Yokota Rizzo

Immunostaining of the clinical samples, microscope pictures acquisitions, and data analysis

### Saskia von Stillfried

Analysis and interpretation of the immunohistochemistry of clinical samples

### Kalle Kilk

polymersome characterization

### Twan Lammers

Critical revision of the manuscript

### Sergio E Moya

Manuscript discussion, edition, and critical revision

### Tambet Teesalu

Manuscript writing, discussion, edition, and critical revision

## Acknowledgements

We acknowledge Prof. Erkki Ruoslahti (Cancer Research Center, Sanford Burnham Prebys Medical Discovery Institute) for critical reading of the manuscript. Rein Laiverik (Department of Anatomy, University of Tartu) and Angel Martinez Villacorta (CIC Biomagune) are acknowledged for their help with the TEM equipment in this work.

